# Repeated co-option of a conserved gene regulatory module underpins the evolution of the crustacean carapace, insect wings and other flat outgrowths

**DOI:** 10.1101/160010

**Authors:** Yasuhiro Shiga, Yasuhiko Kato, Yuko Aragane-Nomura, Takayuki Haraguchi, Theodora Saridaki, Hajime Watanabe, Taisen Iguchi, Hideo Yamagata, Michalis Averof

## Abstract

**Summary statement:** The genes *vestigial, scalloped* and *wingless* comprise a conserved regulatory module that was co-opted repeatedly for the evolution of flat structures, such as insect wings, and crustacean carapace, tergites and coxal plates.

**Summary:** How novelties arise is a key question in evolutionary developmental biology. The crustacean carapace is a novelty that evolved in the early Cambrian. In an extant crustacean, *Daphnia magna*, the carapace grows from the body wall as a double-layered sheet with a specialized margin. We show that the growing margin of this carapace expresses *vestigial, scalloped* and *wingless*, genes that are known to play key roles in regulating growth at the insect wing margin. RNAi-mediated knockdown of *scalloped* and *wingless* impair carapace development, indicating that carapace and wing might share a common mechanism for margin outgrowth. However, carapace and wings arise in different parts of the body and their margins have different orientations, arguing that these structures have independent evolutionary origins. We show that *scalloped* is also expressed at the margin of unrelated flat outgrowths (tergites and coxal plates) in the distantly related crustacean *Parhyale hawaiensis*. Based on these observations, we propose that the *vestigial-scalloped-wingless* gene module has a common role in the margin of diverse flat structures, originating before the divergence of major crustacean lineages and the emergence of insects. Repeated co-option of this module occurred independently in the carapace, wing and other flat outgrowths, underpinning the evolution of distinct novelties in different arthropod lineages.

## Introduction

The carapace, the flat shield that covers the dorsal part of the body in many crustaceans, is an evolutionary novelty that appeared in the early Cambrian, approximately 525 million years ago (Caron and Vannier, 2015; Wills et al., 1998; Xian-guang et al., 2004). The evolutionary origin of the carapace has been the subject of extensive study and debate (Calman, 1909; Dahl, 1991; Fryer, 1996; Walossek, 1993), as the dorsal part of head displays a wide morphological diversity among crustacean lineages. Several questions on the origin of this structure remain unresolved, including whether the carapace is a primitive structure with a single origin or whether it had evolved multiple times, from which segment(s) it originated, and whether it is related to structures in the body of other arthropods. Comparative developmental studies may provide new opportunities to address these questions, by exploring the genetic/developmental mechanisms that underpin the formation of this structure and examining how these mechanisms evolved.

The cladoceran crustacean *Daphnia magna* (Branchiopoda, Cladocera), known as the water flea, is a common inhabitant in lakes and ponds worldwide. *Daphnia* develops a large dorsal carapace (also called a secondary shield, (Walossek, 1995; Walossek, 1993)) that covers the entire trunk region of the adult. Here we describe the development of *Daphnia*’s carapace and identify regulatory genes involved in the growth and patterning of this structure. First, we examine the expression of Hox proteins to address the segmental origin of the carapace. Our study reveals that the *Daphnia* carapace has a cephalic (maxillary) origin. Second, we examine genes involved in the growth and patterning of this structure.

The *Daphnia* carapace is a flat structure, which consists of a double epidermal cell layer decorated with bristles on its margin (Fryer, 1991). These features are also found in other structures in arthropods, including the wings of insects. The development of insect wings has been extensively studied in *Drosophila* and a relatively good understanding of the gene regulatory networks that govern the patterning and growth of insect wings has emerged (Hariharan, 2015). We report that, unexpectedly, some of the regulatory genes that are known to play a key role in patterning the wing and organizing growth around the wing margin – *vestigial (vg), scalloped (sd)* and *wingless (wg)* – have similar roles in the margin of the *Daphnia* carapace.

In *Drosophila, vg* and *sd* are expressed in the wing primordium, where they act as ’selector genes’ to establish the wing fate (Campbell et al., 1992; Guss et al., 2001; Kim et al., 1996; Williams et al., 1991; Williams et al., 1993; Williams et al., 1994); ectopic expression of these genes can induce ectopic wing-like structures (Kim et al., 1996; Paumard-Rigal et al., 1998). *wg* is expressed in a stripe of cells at the prospective wing margin, providing a long-range signal that coordinates growth and patterning around that margin (the ‘wing margin organizer’, (Couso et al., 1994; Diaz-Benjumea and Cohen, 1995; Neumann and Cohen, 1997; Zecca et al., 1996)). Cross-regulatory interactions between *wg, vg* and *sd*, centered around the wing margin, play a crucial role in establishing their expression patterns (Klein and Martinez-Arias, 1999; Williams et al., 1994; Zecca and Struhl, 2007a; Zecca and Struhl, 2007b).

In *Daphnia*, we find that VG, SD and WG proteins are co-expressed at the carapace margin as the carapace develops and extends over the body. RNAi-mediated knockdowns suggest that these genes play a crucial role in carapace formation that extends beyond their expression domain at the margin. These results suggest that a conserved regulatory module comprising *vg, sd* and *wg* is associated with growth around a margin organizer both in *Drosophila* wings and in the *Daphnia* carapace. We propose that this module represents an ancient mechanism that was co-opted independently during arthropod evolution to generate a wide range of flat structures.

## Materials and Methods

### Animals and molecular cloning

Handling of *Daphnia magna* culture and isolation of Hox cDNAs used in this study were described previously (Shiga et al., 2006; Shiga et al., 2002). Degenerate PCR primers used to isolated *D. magna* orthologs of *sd, wg*, and *hh* were 5’-GTBTGCTCHTTYGGCAAGCAAGCARGTGGT-3’ and 5’-AAGTTYTCCAGCACRCTGTTCATCAT-3’, 5’-CGGGATCCATAGAGTCCTGCACCTGCGACTA-3’ and 5’-CGGGATCCGGACATGCCRTGGCATTTGCA-3’, and 5’-CGGGATCGGWGCVGACMGSCTSATG-3’ and 5’-CGGATCCAGTCRAAKCCRGCYTCSAC-3’, respectively. The primers used to isolate *D. magna vg, ap1*, and *ap2*, based on the *vg*- and *ap*-related sequences found in a draft genome sequences of *D. pulex*, were 5-CAGTACATTTCAGCTAATTGCGTCGT-3’ and 5’-TTGAGGGCCCGGCTGAAATGCTGGTC-3’, 5’-TCGCCGTCTCCGACAACTCTGTG-3’ and 5’-GCCATCTTGTTGTCGCATTATGTTGCG-3’, and 5’-AGCGAGTGTCTCAGGTGCGACGAA-3’ and 5’-CTTGGAGAGGGTGGTCTTCTGCGA-3’. Using *D. magna* cDNA library as a template, PCR products were amplified, cloned into pCR4-Topo (Invitrogen) or pUC118, and used as probes to screen full-length cDNA clones. The *sd* ortholog of *Parhyale hawaiensis* was isolated by nested PCR on cDNA from mixed embryonic stages, using degenerate primers 5’-CGGGATCCGARCARAGYTTYCARGA-3’, 5’-GGAATTCGAYGARGGIAARATGTA-3’ and 5’-GCTCTAGAACICTRTTCATCATRTA-3’. *D. magna* sequences have been deposited in the GenBank/DDBJ/EMBL database under the following accession numbers: *Scr*, AB465513; *Dfd*, AB539164; *vg*, AB465512; *sd* (short form), AB465514; *sd* (long form), AB465515; *wg*, AB465516; *hh*, AB465517; *ap1*, AB539165; *ap2*, AB539166. *Parhyale hawaiensis* sequence for *sd* was deposited under FN256248. *D. pulex* sequence data were produced by the US Department of Energy Joint Genome Institute (http://www.jgi.doe.gov/) in collaboration with the *Daphnia* Genomics Consortium (http://daphnia.cgb.indiana.edu).

### Immunostaining and whole mount in situ hybridization

Rat anti-SCR, rat anti-SD, rabbit anti-VG, and rabbit anti-DFD were raised against recombinant proteins (residues 1-125, 1-475, 1-381, and 1-296, respectively) and affinity purified. For VG, a rabbit antibody was also raised against the synthetic peptide GLEAGQVQQEPGKDLYWF, corresponding to the C-terminal sequence of VG (anti-VG-pep). Both VG antibodies gave identical results. Rabbit anti-ANTP, rat anti-UBX, and monoclonal anti-PDM/NUB Mab 2D4 were described previously (Damen et al., 2002; Shiga et al., 2002; Shiga et al., 2006). Engrailed expression was visualized using the monoclonal antibody 4F11 (a gift from Dr. Nipam Patel) (Patel et al., 1989b). Methods for immunostaining and in situ hybridization for whole-mount *Daphnia* embryos (Shiga et al., 2002) are available upon request. Embryos were classified according to the staging system for *D. magna* embryogenesis (Mittmann et al., 2014). Stained embryos were observed under a Leica TCS-SP2 (Solms, Germany) or Olympus FV-1000 (Tokyo, Japan) confocal microscopes. In situ hybridization and staging of *Parhyale hawaiensis* embryos were carried out as described previously (Rehm et al., 2009).

### RNA interference

*D. magna* RNAi experiments were performed according to the protocol by Kato *et al*. (Kato et al., 2011). Double-strand RNAs (dsRNAs) were synthesized from portions of *vg, sd*, and *wg* cDNAs with MEGAscript T7 Kit (Ambion) and approximately 0.3 nl of each dsRNA solution, at concentrations of 37, 111, 333, and 1000 μg/ml, were microinjected into *D. magna* embryos at the earliest stage. As controls, we injected *D. magna white* and *E. coli malE* dsRNAs at 1000 μg/ml, were used. Injected embryos were fixed after 45 hours of cultivation at 24°C, stained with the nuclear dye YOYO-1, and observed for carapace phenotypes. For double immunostaining of *wg* RNAi embryos, samples were fixed after 35 hours. To exclude the possibilities of non-specific off-target effects, two dsRNAs synthesized from non-overlapping regions of each cDNA were tested: *sd* #1, 689 bp in length corresponding to the central portion of SD containing the C-teriminal region of the TEA domain through the N-terminal region of the VBD; *sd* #2, 683 bp corresponding to the C-terminal region of the VBD plus the 3’-UTR; *wg* #1, 683 bp corresponding to the central portion of the mature WG; *wg* #2, 611 bp corresponding to the 5’-UTR and the N-terminal region of WG including the signal peptide.

## Results and Discussion

### The *Daphnia* carapace is a double-layered sheet that grows from maxillary segments

Carapace development in *Daphnia* involves the outgrowth of a double-layered epithelial sheet with a discrete margin (Figure 1). As the outgrowth expands posteriorly and laterally, the inner and outer layers of the carapace become folded onto each other. At the edge where the inner and outer layers meet there is a morphologically distinct margin. In if posterior-most region, and the midline, forms a long and narrow projection called the apical spine (or carapace spine) (Fryer, 1991; Kotov and Boikova, 2001) (Figure 1). The mature carapace comprises a double epidermal layer decorated with several types of bristles on its margin (Fryer, 1991) (Figures 1I and 1J).

**Figure 1.**
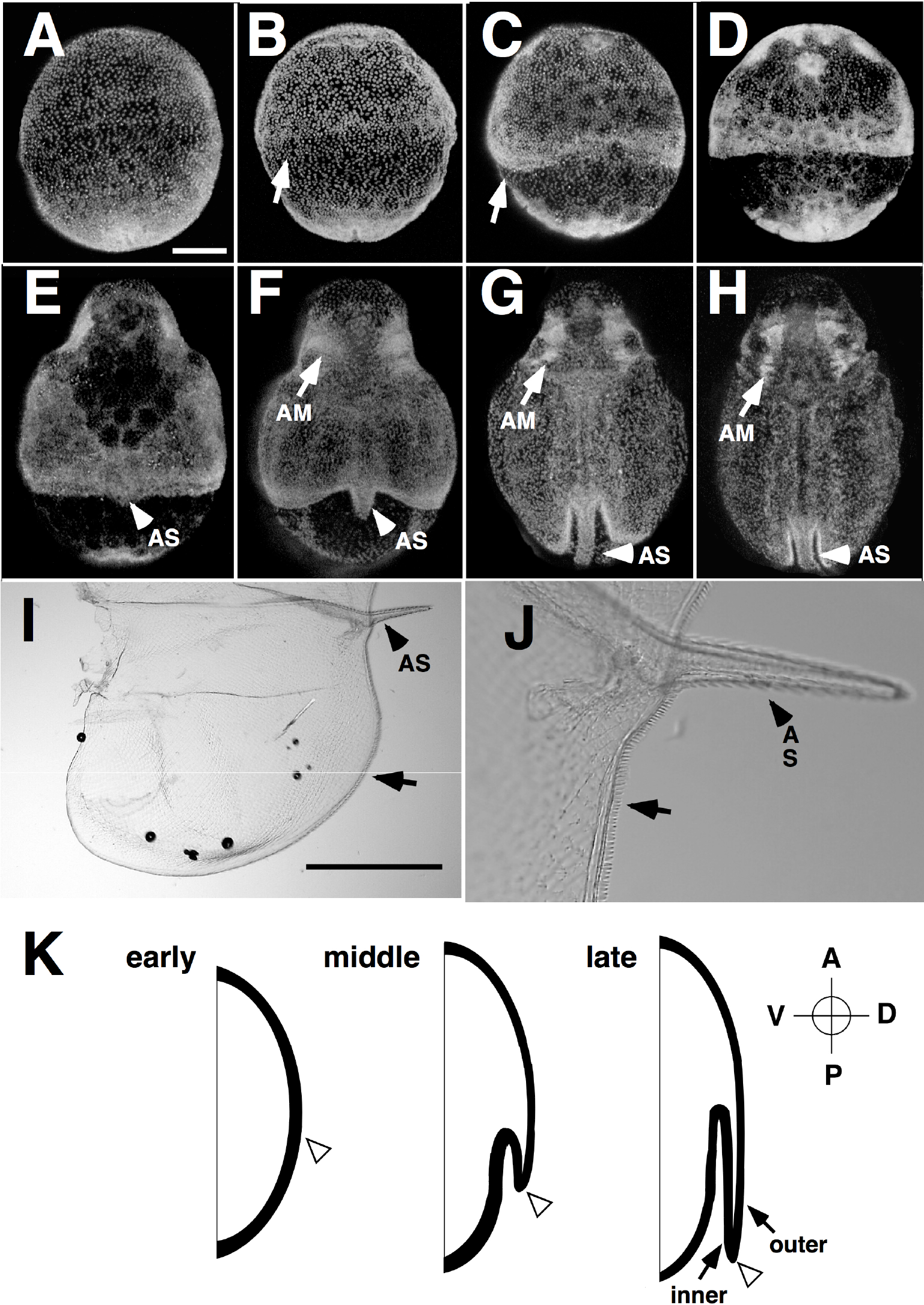
*Daphnia* carapace development. **(A-H)** Carapace development for *D. magna* embryos cultured at 24°C. Embryos were fixed, stained with the nuclear dye YOYO-1, and imaged from the dorsal side. (A) Stage 7.1, stage just prior to appearance of carapace primordium; (B) Stage 7.5, slight crowding of nuclei in the lateral and dorsal regions; (C) early Stage 8, rudiments of the carapace appear as epithelial folds on the lateral side of the embryos; (D) late Stage 8, progression of carapace folding is significantly faster on the lateral side than in the dorsal-most region; (E) Stage 9, apical spine (AS) starts to elongate; (F-H) Stages 10, 11 and 12, posterior and lateral carapace extension and apical spine elongation continue. Antennal muscles (AM) appear and develop. (H) Carapace covers almost the entire trunk region. (**I** and **J**) Morphology of adult *Daphnia* carapace. Dissected half of the bivalved carapace (anterior left, dorsal up), with apical spine (arrowhead) and several types of marginal bristles (arrow). (**K**) Schematic representation of carapace outgrowth relative to the anteroposterior (AP) and dorsoventral (DV) axes of the body. The *Daphnia* carapace develops as an epithelial fold with a distinct margin (arrowhead) that extends posteriorly and covers the entire trunk region. Scale bars: (A) 100 μm and (I) 1 mm.

To determine the segmental origin of the carapace, we examined the expression of ten Hox proteins present in the *Daphnia* genome ((Shiga et al., 2002; Shiga et al., 2006) and unpublished). Sex Combs Reduced (SCR) was the only Hox protein expressed in the carapace (Figure 2) throughout its development, displaying clear boundaries anteriorly and posteriorly with the expression domains of Deformed (DFD) and Antennapedia (ANTP), respectively (Figures 2J-2R). Carapace outgrowth started in a narrow band of cells with reduced levels of SCR, within the SCR expression domain (Figures 2E, 2N, and 2O).

**Figure 2.**
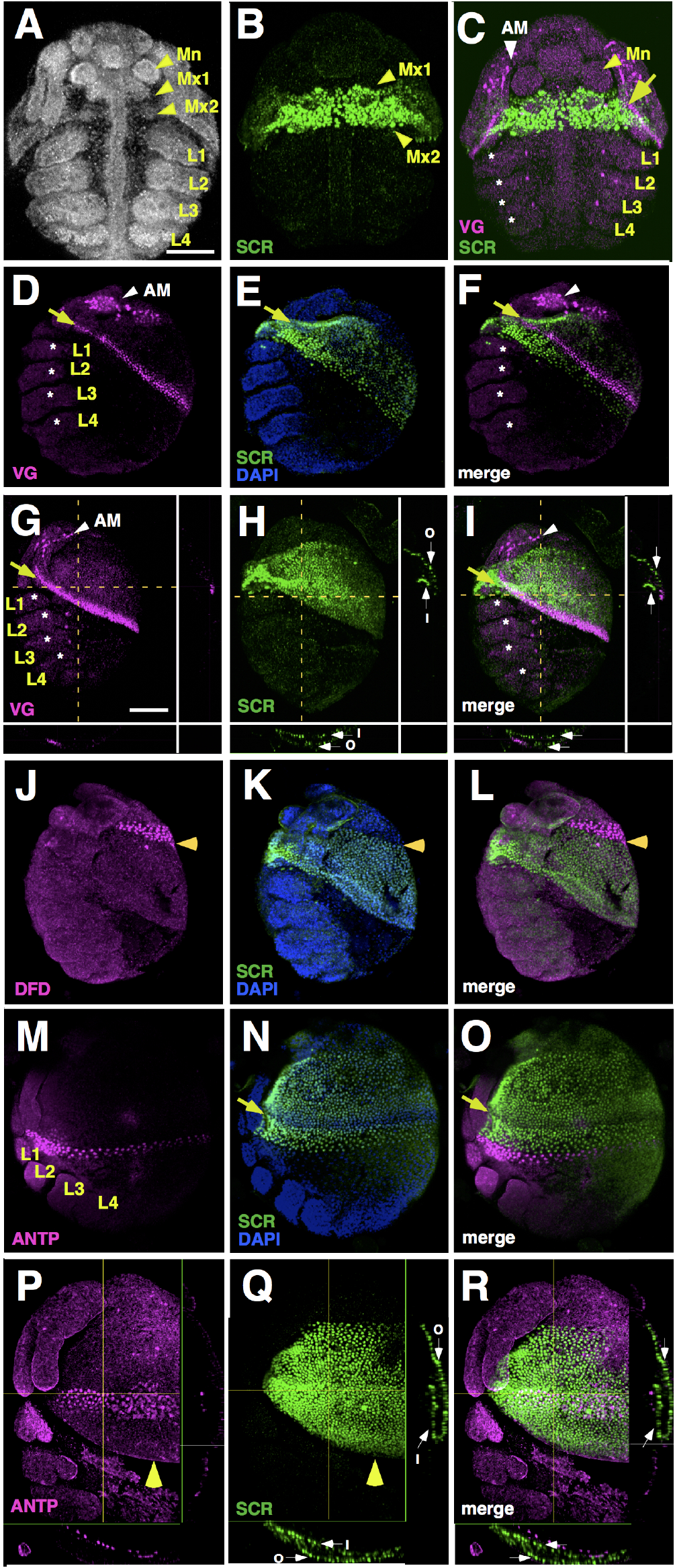
Expression of SCR and VG proteins in developing *Daphnia* embryos. **(A-C)** Ventral view of embryos at Stage 8, stained with the nuclear dye YOYO-1 (A) or with anti-SCR and anti-VG (B and C). Ventral SCR expression is restricted within the Mx1 and Mx2 segments. (**D-I**) Lateral views of embryos double stained with anti-SCR (green) and anti-VG (magenta), at Stage 7.4 (D-F) and Stage 8 (G-I). Images of coronal and transverse confocal sections are shown on the lower and right sides of panels G-I, respectively. VG expression in the carapace margin lies entirely within the SCR domain (yellow arrows). Prior to carapace folding (D-F), dorsal SCR expression is divided into a cell-crowded anterior and a less crowded posterior, by a belt of cells with low SCR expression, in which VG expression is activated (arrows). After carapace folding begins (G-I), SCR is expressed in both the outer and the inner epithelium of the developing carapace (white arrows marked O and I, respectively). The epipodites (asterisks) lack VG expression. (**J-L**) Lateral views of an embryo at Stage 8, stained with anti-SCR (green), anti-DFD (magenta) and the nuclear dye DAPI (blue). SCR-expressing domain displays a clear anterior boundary (horizontal arrowheads) with the expression domain of DFD. (**M-R**) Lateral view of embryos with anti-SCR (green), anti-ANTP (magenta) and the nuclear dye DAPI (blue), at Stage 7.4 (M-O) and Stage 9 (P-R). Coronal and transverse confocal sections are shown on the lower and right sides of panels P-R, respectively. SCR and ANTP are expressed in adjacent but mutually exclusive territories in the dorsolateral epidermis (M-O). The low SCR expressing belt where VG expression is activated is clearly visible (yellow arrows in N and O). As the carapace grows, it extends over the ANTP-expressing domain (growing carapace margin marked by yellow arrowheads, P-Q). Mn, mandible; Mx1, first maxillae; Mx2, second maxillae; L1 to L4, first to fourth thoracic legs. Scale bars: 100 pm. See also Figures S1.

In *Daphnia*, SCR is expressed in the first and second maxillary segments (M×1 and M×2, respectively), strongly suggesting that the *Daphnia* carapace is derived from the M×1 and/or the M×2 segments (Figures 2A-2C). This result corroborates previous observations tracing the origin of the cladoceran (anomopodan) carapace to the M×2 segment (Fryer, 1991; Fryer, 1996; Kotov and Boikova, 2001).

### *vestigial, scalloped* and *wingless* are expressed at the growing carapace margin

To determine the expression patterns of *Daphnia* VG and SD we raised specific antibodies against these proteins and we performed antibody stainings in *Daphnia* embryos. The expression patterns that we observed were also confirmed by in situ hybridization (data not shown). The first detectable VG expression is observed in the developing large muscles of the second antenna, around Stage 7.2. Around Stage 7.4, VG expression appears in the prospective carapace, in the band of cells with reduced SCR expression (Figures 2D-2F). VG expression first appears in the lateral borders of the carapace primordium and then extends dorsally (Stages 7.5 to 8, Figures S1I and S1J). Folding of the carapace follows VG expression, initiating laterally (Figures S1I and S1M) and then extending to the dorsal-most region (Figures S1J and S1N). The apical spine starts to develop in Stage 9 (Figure 1E). VG expression is initially widespread in the elongating apical spine (Stages 9 to 10, Figure 3B; Figures S1K and S1O) and then becomes restricted to the center of the apical spine (Figures S1D and S1H). SD expression was observed in the developing carapace margin and apical spine (Figures 3A and 3C; Figures S1A-S1H), as well as in the antennal muscles, labrum, mandible (Mn), Mx1, and the region around the anus (data not shown). SD is co-expressed with VG in the carapace and antennal muscles (Figures 3A-3C), consistent with the cooperative action established for these two proteins in *Drosophila* (Halder and Carroll, 2001; Halder et al., 1998; Paumard-Rigal et al., 1998; Simmonds et al., 1998).

**Figure 3.**
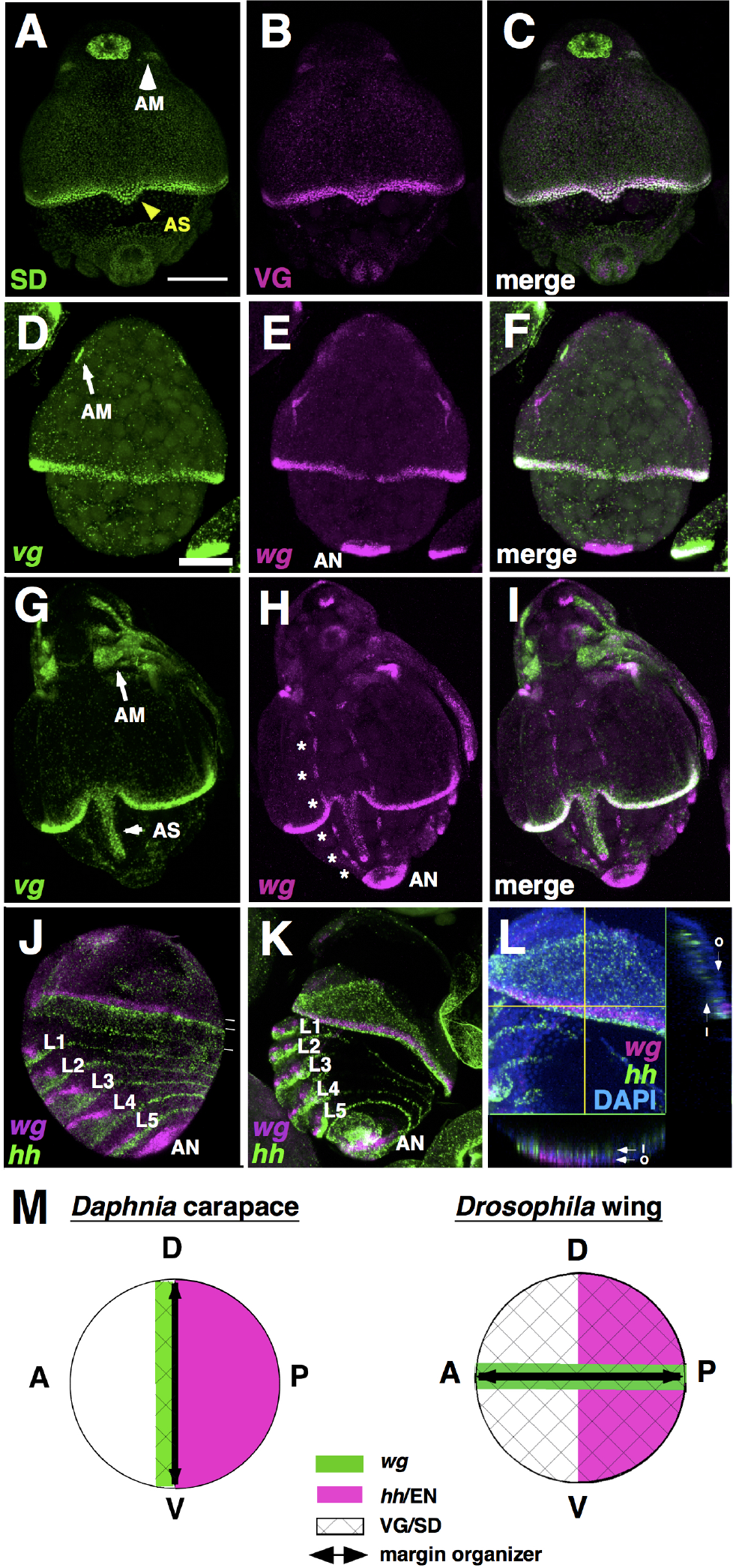
Expression of VG, SD and WG at the *Daphnia* carapace margin. **(A-C)** Dorsal view of an embryo at Stage 9, double stained with anti-SD (green) and anti-VG (magenta). VG and SD are co-expressed in cells of the developing carapace margin, apical spine (AS) and antennal muscles (AM). **(D-I)** Dorsal view of embryos in which expression of *vg* (green) and *wg* (magenta) is visualized by double in situ hybridization, at Stage 8 (D-F) and Stage 10 (G-I). Co-expression of *vg* and *wg* is evident in the carapace margin and apical spine. Note that widths of *vg*- and wg-expressing domains in the carapace margin are the same. *vg* is also expressed in antennal muscles (arrows in D and G), while *wg* is also detected in the thoracic legs (arrowheads), anal region (AN) and trunk (asterisks). (**J-L**) Lateral views of embryos in which expression of *hh* (green) and *wg* (magenta) is visualized by double in situ hybridization, at Stage 7.3 (J) and Stage 8 (K and L); nuclei are stained with DAPI (blue). As in all arthropod species studied to date, *wg* and *hh* are expressed in adjacent territories, within the anterior and posterior compartment of segments, respectively. Unlike *hh* expression, *wg* expression is restricted to the ventral part of each segment (up to the dorsal limit of thoracic appendages L1-L5), except in the regions of the anus and the presumptive carapace margin, where the neighboring territories of *hh* and *wg* extend to the dorsal side (J). The widths of the *wg*- and *hh*-expressing domains at the dorsal side of carapace anlage are indicated by white bars. As the carapace fold grows (K and L), *wg* is expressed at the posterior border of of the outer layer of the carapace, while *hh* is expressed widely in the inner layer (marked O and I, respectively, in the optical sections in L). (**M**) Schematic representation of gene expression patterns around the margins of the *Daphnia* carapace and the *Drosophila* wing. The margin (black bar) appears where the expression of *wg* (green) and VG/SD (mesh) overlap. In *Daphnia*, the carapace margin forms at the anteroposterior (AP) compartment boundary, between the expression domains of *wg* and EN (magenta), while in *Drosophila* wings it lies on the dorsoventral (DV) compartment boundary, which lies perpendicular to the AP compartment boundary. Scale bar: 100 μm. See also Figures S1, S2, and S3.

Double in situ hybridization also revealed the broadly overlapping expression of *wg* and *vg* in the outer epithelial layer of the carapace margin of *Daphnia* throughout its development (Figures 3D-3I; Figures S2A-S2C). Unlike *Drosophila* wings, where the expression of VG and SD extends far beyond the wing margin, driven by an autoregulatory loop and the long-range diffusion of WG (Neumann and Cohen, 1997; Zecca and Struhl, 2007a; Zecca et al., 1996), the expression domains of WG, VG and SD fully overlap in the *Daphnia* carapace margin (Figures 3A-3I; Figures S2A-S2C).

In *Drosophila*, the expression of *wg* and the establishment of an organizer at the wing margin depend on the *apterous (ap)* gene, which is expressed in the dorsal compartment of wing discs (Diaz-Benjumea and Cohen, 1993; Diaz-Benjumea and Cohen, 1995). The localized expression of *apterous* directs the activation of Notch signaling at the margin, which in turn activates the expression of *wg*. The *Daphnia* genome contains two *ap* orthologues, *ap-1* and *ap-2*. We cloned these and studied their expression. *ap1* is expressed in the epipodites and in the nervous system, but not in the developing carapace (Figures S3A-S3F). *ap2* is expressed in the epipodites, the nervous system, the labrum and in the carapace margin (Figures S3G-S3L), However, this latter expression at the margin appears during late stages of carapace elongation, long after the onset of WG, VG and SD expression, (Figures S3I, S3J, and S3M-S3O). We therefore do not think that *apterous* in involved in setting up *wg* expression at the *Daphnia* carapace margin.

We also examined the expression of Notch signaling pathway (NSP) components, including *Notch, Serrate, Delta* and *fringe* genes, by in situ hybridization. We found no expression of these genes associated with the early stages of carapace margin outgrowth. Furthermore, knockdown of NSP ligands *Delta* and *Serrate* did not affect carapace specification and elongation (Uehara and Shiga, unpublished observations). These data indicate that the *Daphnia* carapace and *Drosophila* wing share a common set of genes that are expressed during margin outgrowth, which include VG, SD and WG, but not AP or components of the NSP.

### RNAi reveals essential functions for carapace outgrowth

To directly test the functions of *vg, sd*, and *wg* in *Daphnia* carapace development, we knocked these genes down by RNA interference (RNAi). Although the injection of high amounts of dsRNA was lethal for all three genes (Table S1), *sd* and *wg* RNAi embryos developed to late stages, permitting phenotypic analyses at 45 hours AED, corresponding to Stage 12. To ensure that the observed phenotypes were not due to off-target effects, two dsRNAs synthesized from non-overlapping regions of the corresponding cDNAs were tested for each gene (Table S2). In mildly affected *sd* RNAi embryos, the carapace margin was greatly reduced (Figure 4B) and the apical spine was shorter than that in wild type embryos, or in embryos injected with control dsRNAs (Figure 1G; Figure 4A; Table S1). In more severe cases, carapace formation was strongly affected (Figure 4C) and in extreme cases only a trace of the carapace remained (Figure 4D). Knockdown of *wg* often resulted in reduction of the trunk region (Figures 4E-4G) and malformation of ventral appendages (Figures 4H and 4I). In mildly affected *wg* RNAi embryos, the carapace was heavily wrinkled and the apical spine was malformed (Figure 4E). In more severe cases, carapace and apical spine outgrowth were severely impaired (Figure 4F) and the carapace fold was barely observable (Figure 4G), except for small regions around the putative carapace margin (Figure 4H). These experiments demonstrate that SD and WG play essential roles in carapace outgrowth and in the elongation of the apical spine. The phenotypes extend beyond the expression domains of SD and WG at the carapace margin, with non-autonomous effects on the growth of the entire carapace, suggesting that the carapace margin may have organizer properties analogous to those of the wing margin in insects.

**Figure 4.**
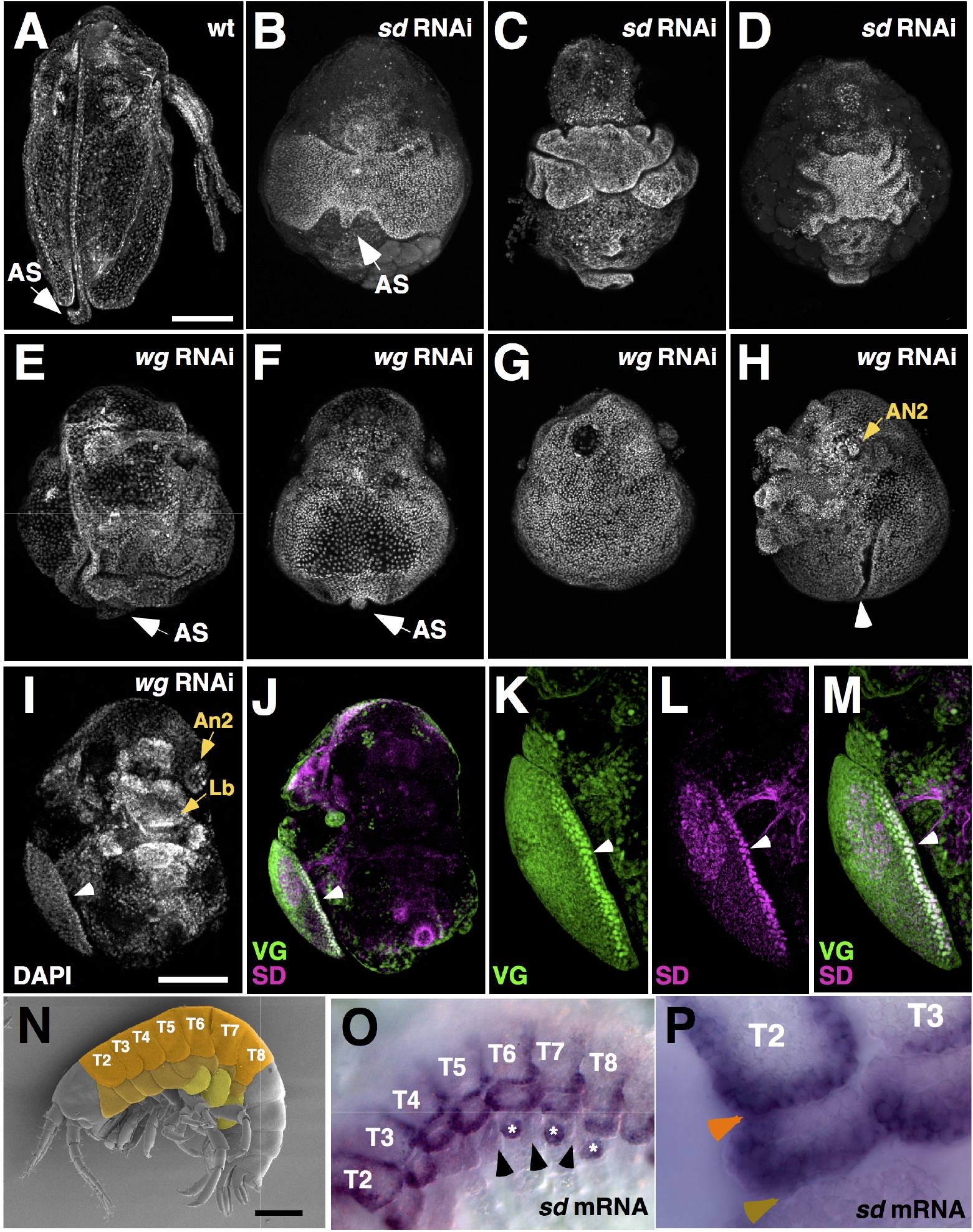
Functional evaluation of *sd* and *wg* in *Daphnia* and association with other flat outgrowths. (**A**) Dorsal view of a wild-type (mock-injected) embryo at 45 hours after egg deposition (AED), corresponding to Stage 12. (**B-G**) Dorsal view of embryos at 45 hours AED, microinjected with 300 pg of *sd* (B-D) or *wg* (E-G) dsRNAs, showing typical ‘mild’ (B and E), ‘medium’ (C and F), and ‘strong’ (D and G) carapace phenotypes. (**H**) Lateral view of the same embryo as in (G). (B) Carapace outgrowth and apical spine elongation are impaired. (C) The carapace is extensively disintegrated. (D) Only a trace of the carapace remains. (E) Heavily wrinkled carapace and curled apical spine are observed. (F) Carapace outgrowth is severely impaired and apical spine is stunted. (G and H) Carapace fold is barely observable, except for small regions around the probable carapace margin (arrowhead); ventral appendages are heavily distorted (on the left in H). (**I-M**) Ventral views of an embryo at 35 hours AED, corresponding to Stage 9, microinjected with 300 pg of *wg* dsRNA and stained with anti-VG (green), anti-SD (magenta) and the nuclear dye DAPI (shown in I). The embryo has heavily distorted appendages and an unexpanded carapace, but VG and SD are co-expressed at carapace margin (white arrowhead) as in wild type embryos (see Figure 3C). (**N**) Electron micrograph of *Parhyale hawaiensis* hatchling, highlighting the morphology of tergites (orange), coxal plates on T2 to T8 (brown), and flat outgrowths on the basis of thoracic appendages T6 to T8 (yellow). (**O** and **P**) In situ hybridization for *sd*in stage 23-24 *Parhyale* embryo. Expression is seen on the margin of tergites, coxal plates and flat outgrowths on the basis of appendages T6-T8 (asterisks). No expression is seen in the epipodites/gills (arrowheads). Higher magnification of the T2 appendage (P) shows expression in the margin of tergites, and coxal plates, marked by orange and brown arrowheads, respectively. Scale bars: (A,I) 100 μm and (N) 200 μm. See also Tables S1 and S2.

In *Drosophila, wg* function is essential for initiating and maintaining the expression of VG in the wing primordium (Klein and Martinez-Arias, 1999; Zecca and Struhl, 2007a; Zecca and Struhl, 2007b). To test whether VG or SD expression is affected by *wg* knockdown in *Daphnia*, we carried out double immunostainings for VG and SD in *wg* RNAi embryos. We found that both proteins are expressed, even in embryos with a strong phenotype (Figures 4J-4M), indicating that high levels of *wg* expression are not necessary for VG and SD expression at the carapace margin.

### The VG-SD-WG module was independently co-opted in carapace, wings and other flat outgrowths

Does a shared set of genes regulating insect wing and *Daphnia* carapace development mean that these structures are homologous? The orientation of wing and carapace with respect to the anterior-posterior (A/P) and dorsal-ventral (D/V) body axes argues strongly against this notion. In all arthropods, *engrailed (en)* and *hedgehog (hh)* are consistently expressed at the posterior compartment of each segment, encompassing the posterior part of appendages such as legs and wings (Basler and Struhl, 1994; Damen, 2002; Patel et al., 1989a). The insect wing margin forms along a D/V boundary that runs perpendicular to the segmental stripes of *en/hh* expression (Basler and Struhl, 1994; Diaz-Benjumea and Cohen, 1993). In contrast, we find that the *Daphnia* carapace margin is established along the A/P compartment boundary: the *Daphnia* orthologs of *en* and *hh* are expressed in the posterior part of the carapace, bordering the VG-, SD-, and *wg*-expressing domains at the carapace margin (Figure 3J; Figures S2G-S2I). Their expression persists in the internal layer of carapace after the initiation of carapace folding (Figures 3K and 3L; Figures S2D-S2F and S2J-S2L). This fundamental difference in the topology of the insect wing and the *Daphnia* carapace, relative to a highly conserved A/P landmark (Figure 3M), suggests that the two structures have co-opted the VG, SD and WG independently at their margin.

To address whether this gene module has been recruited in other structures during crustacean evolution, we cloned and studied the expression of an *sd* ortholog in the amphipod *Parhyale hawaiensis* (Malacostraca: Amphipoda). *Daphnia* and *Parhyale* belong to two major divergent clades of crustaceans, branchiopods and malacostracans, respectively, which span a large evolutionary distance. *Parhyale* develops no carapace on its dorsal side. Strikingly, *sd* in *Parhyale* is specifically expressed at the margin of the tergites, the flat ectodermal plates that cover the dorsal part of each trunk segment (Figures 4N-4P). Expression is also prominent in the flat coxal plates of thoracic appendages T2 to T8, and in flat outgrowths at the bases of thoracic appendages T6 to T8 (Figures 4N-4O). This expression is observed during the embryonic stages when these structures are growing (Figure 4P), reinforcing the strong association of *sd* expression and the outgrowth of flat structures in different parts of the body. Based on these observations, we propose that a molecular program for sheet-like outgrowth, mediated by VG, SD and WG, was established before the divergence of major crustacean lineages and the emergence of insects. Our data suggest that this programme was co-opted independently to pattern diverse flat structures in different crustaceans and insect lineages. This hypothesis is consistent with observations on *vg* gene expression and function in diverse body wall outgrowths in insects (Clark-Hachtel et al., 2013; Niwa et al., 2010; Ohde et al., 2013).

### Implications for the origin of insect wings

Previous studies have raised two contrasting hypotheses on the origin of insect wings: one suggests that wings evolved as novel extensions of the body wall (Snodgrass, 1935); the second holds that wings evolved from pre-existing dorsal appendage branches, epipodites, that usually function as gills in aquatic arthropods (Wigglesworth, 1973). The second hypothesis is supported by the observation that the transcription factors Nubbin (NUB) and Apterous (AP) are expressed in the wings of insects and in the epipodites of crustaceans (Averof and Cohen, 1997) (see Figure S3). However, the epipodites of *Daphnia* and *Parhyale* do not express VG, SD or *wg* (asterisks in Figure 2 and arrowheads in Figure 4O) and both lack a sheet-like structure with a distinct margin. More recent studies have argued that wing primordia could have a dual origin, deriving from the fusion of body wall and limb elements (Clark-Hachtel et al., 2013; Elias-Neto and Belles, 2016; Niwa et al., 2010; Ohde et al., 2013).

Our results suggest an alternative way to reconcile these hypotheses: we propose that wings evolved by integrating developmental modules that were previously active in different parts of the body, namely, by co-option of the VG-SD-WG module, that was associated with flat body wall outgrowths, onto epipodites expressing AP and NUB. Placing the VG-SD-WG module under the control of a stable D/V boundary via AP may have been a key step in assembling the gene regulatory network of the wing.

Generating novelty by combining pre-existing elements may be a common feature of evolution, an idea captured in the notion that ‘evolution is a tinkerer’ (Jacob, 1977). It is striking that independent co-option of the same regulatory module could produce flat structures with such important and diverse roles in crustacean and insect lineages.

## Acknowledgements

We thank Drs. S. Hayashi and N. Niwa for sharing unpublished observations and critical reading of the manuscript, Dr. S. Tokishita for assistance in phylogenetic analyses, H. Sato and A. Akimoto-Kato for technical assistance on molecular cloning and antibody preparation, and Dr. N. Patel for the 4F11 monoclonal antibody. This work was supported by a grant from the Ministry of Education, Culture, Sports, Science and Technology of Japan to Y.S. and grants from the Ministry of Environment, Japan and Long-range Research Initiative by Japan Chemical Industry Association to T.I.. Y.K. was a Research Fellow of the Japan Society for the Promotion of Science.

## References

Averof, M., and Cohen, S. M. (1997). Evolutionary origin of insect wings from ancestral gills. Nature 385, 627–630.

Basler, K. and Struhl, G. (1994). Compartment boundaries and the control of Drosophila limb pattern by hedgehog protein. Nature 368, 208–214.

Calman, W. T. (1909). A Treatise on Zoology; Crustacea. (ed. Lankester, R. London: Adam and Charles Black.

Campbell, S., Inamdar, M., Rodrigues, V., Raghavan, V., Palazzolo, M. and Chovnick, A. (1992). The scalloped gene encodes a novel, evolutionarily conserved transcription factor required for sensory organ differentiation in Drosophila. Genes & Development 6, 367–379.

Caron, J.-B. and Vannier, J. (2015). Waptia and the Diversification of Brood Care in Early Arthropods. Curr. Biol.

Clark-Hachtel, C. M., Linz, D. M. and Tomoyasu, Y. (2013). Insights into insect wing origin provided by functional analysis of vestigial in the red flour beetle, Tribolium castaneum. Proc. Natl. Acad. Sci. U.S.A.

Couso, J. P., Bishop, S. A. and Martinez-Arias, A. (1994). The wingless signalling pathway and the patterning of the wing margin in Drosophila. Development 120, 621–636.

Dahl, E. (1991). Crustacea Phyllopoda and Malacostraca: a Reappraisal of Cephalic and Thoracic Shield and Fold Systems and Their Evolutionary Significance. Philos. Trans. R. Soc. Lond., B, Biol. Sci. 334, 1–26.

Damen, W. (2002). Parasegmental organization of the spider embryo implies that the parasegment is an evolutionary conserved entity in arthropod embryogenesis. Development 129, 1239–1250.

Damen, W.G.M., Saridaki, T. and Averof, M. (2002). Diverse adaptations of an ancestral gill: a common evolutionary origin for wings, breathing organs, and spinnerets. Curr. Biol. 12, 1711–1716.

Diaz-Benjumea, F. J. and Cohen, S. M. (1993). Interaction between dorsal and ventral cells in the imaginal disc directs wing development in Drosophila. Cell 75, 741–752.

Diaz-Benjumea, F. J. and Cohen, S. M. (1995). Serrate signals through Notch to establish a Wingless-dependent organizer at the dorsal/ventral compartment boundary of the Drosophila wing. Development 121, 4215–4225.

Elias-Neto, M. and Belles, X. (2016). Tergal and pleural structures contribute to the formation of ectopic prothoracic wings in cockroaches. R Soc. open sci. 3, 160347.

Fryer, G. (1991). Functional Morphology and the Adaptive Radiation of the Daphniidae (Branchipoda, Anomopoda). Philos. Trans. R Soc. Lond., B, Biol. Sci. 331, 1–99.

Fryer, G. (1996). The carapace of the branchiopod Crustacea. Philos. Trans. R. Soc. Lond., B, Biol. Sci. 351, 1703–1712.

Guss, K. A., Nelson, C. E., Hudson, A., Kraus, M. E. and Carroll, S. B. (2001). Control of a genetic regulatory network by a selector gene. Science 292, 1164–1167.

Halder, G. and Carroll, S. B. (2001). Binding of the Vestigial co-factor switches the DNA-target selectivity of the Scalloped selector protein. Development 128, 3295–3305.

Halder, G., Polaczyk, P., Kraus, M. E., Hudson, A., Kim, J., Laughon, A. and Carroll, S. (1998). The Vestigial and Scalloped proteins act together to directly regulate wing-specific gene expression in Drosophila. Genesand Development. 12, 3900–3909.

Hariharan, I. K. (2015). Organ Size Control: Lessons from Drosophila. Dev. Cell 34, 255–265.

Jacob, F.(1977). Evolution and tinkering. Science 196, 1161–1166.

Kato, Y., Shiga, Y., Kobayashi, K., Tokishita, S.-I., Yamagata, H., Iguchi, T. and Watanabe, H. (2011). Development of an RNA interference method in the cladoceran crustacean Daphnia magna. Dev. Genes Evol. 220, 337–345.

Kim, J., Sebring, A., Esch, J. J., Kraus, M. E., Vorwerk, K., Magee, J. and Carroll, S.B. (1996). Integration of positional signals and regulation of wing formation and identity by Drosophila vestigial gene. Nature 382, 133–138.

Klein, T. and Martinez-Arias, A. (1999). The vestigial gene product provides a molecular context for the interpretation of signals during the development of the wing in Drosophila. Development 126, 913–925.

Kotov, A. A. and Boikova, O. S. (2001). Study of the late embryogenesis of Daphnia (Anomopoda, “Cladocera,” Branchiopoda) and a comparison of development in Anomopoda and Ctenopoda. Hydrobiologia 442, 127–143.

Mittmann, B., Ungerer, P., Klann, M., Stollewerk, A. and Wolff, C. (2014) Development and staging of the water flea Daphnia magna (Straus, 1820; Cladocera, Daphniidae) based on morphological landmarks. Evolution & Development 5: 12.

Neumann, C. J. and Cohen, S. M. (1997). Long-range action of Wingless organizes the dorsal-ventral axis of the Drosophila wing. Development 124, 871–880.

Niwa, N., Akimoto-Kato, A., Niimi, T., Tojo, K., Machida, R. and Hayashi, S. (2010). Evolutionary origin of the insect wing via integration of two developmental modules. Evolution & Development 12, 168–176.

Ohde, T., Yaginuma, T. and Niimi, T. (2013). Insect Morphological Diversification Through the Modification of Wing Serial Homologs. Science 340, 495–498.

Patel, N. H., Kornberg, T. B. and Goodman, C. S. (1989a). Expression of Engrailed During Segmentation in Grasshopper and Crayfish. Development 107, 201–212.

Patel, N. H., Martin-Blanco, E., Coleman, K. G., Poole, S. J., Ellis, M. C., Kornberg, T. B. and Goodman, C. S. (1989b). Expression of engrailed proteins in arthropods, annelids, and chordates. Cell 58, 955–968.

Paumard-Rigal, S., Zider, A., Vaudin, P. and Silber, J. (1998). Specific interactions between vestigial and scalloped are required to promote wing tissue proliferation in Drosophila melanogaster. Dev. Genes Evol. 208, 440–446.

Rehm, E. J., Hannibal, R. L., Chaw, R. C., Vargas-Vila, M. A. and Patel, N. H. (2009). In situ hybridization of labeled RNA probes to fixed Parhyale hawaiensis embryos. Cold Spring Harb Protoc 2009, pdb.prot5130–pdb.prot5130.

Shiga, Y., Sagawa, K. and Takai, R. (2006). Transcriptional readthrough of Hox genes Ubx and Antp and their divergent post-transcriptional control during crustacean evolution. Evolution & Development 8, 407–414.

Shiga, Y., Yasumoto, R., Yamagata, H. and Hayashi, S. (2002). Evolving role of Antennapedia protein in arthropod limb patterning. Development 129, 3555–3561.

Simmonds, A. J., Liu, X. F., Soanes, K. H., Krause, H. M., Irvine, K. D. and Bell, J.B. (1998). Molecular interactions between Vestigial and Scalloped promote wing formation in Drosophila. Genes & Development 12, 3815–3820.

Snodgrass, R. E. (1935). Principles of Insect Morphology. In Principles of Insect Morphology, pp. 83–99. New York: McGraw-Hill.

Walossek, D. (1995). The Upper Cambrian Rehbachiella, Its Larval Development, Morphology and Significance for the Phylogeny of Branchiopoda and Crustacea. Hydrobiologia 298, 1–13.

Walossek, D.(1993). The upper Cambrian Rehbachiella and the phylogeny of Branchiopoda and Crustacea. Fossils & Strata 32, 1–202.

Wigglesworth, V. B. (1973). Evolution of insect wings and flight. Nature.

Williams, J. A., Bell, J. B. and Carroll, S. B. (1991). Control of Drosophila wing and haltere development by the nuclear vestigial gene product. Genesand Development. 5, 2481–2495.

Williams, J. A., Paddock, S. W. and Carroll, S. B. (1993). Pattern formation in a secondary field: a hierarchy of regulatory genes subdivides the developing Drosophila wing disc into discrete subregions. Development 117, 571–584.

Williams, J. A., Paddock, S. W., Vorwerk, K. and Carroll, S.B. (1994). Organization of wing formation and induction of a wing-patterning gene at the dorsal/ventral compartment boundary. Nature 368, 299–305.

Wills, M. A., Briggs, D., Fortey, R. A. and Wilkinson, M. (1998). An arthropod phylogeny based on fossil and recent taxa. Arthropod fossils and ….

Xian-guang, H., Aldridge, R. J., Bergström, J., Siveter, D. J. and Xiang-Hong, F. (2004). The Cambrian Fossils of Chengjiang, China: the Flowering of Early Animal Life. Oxford: Blackwell Publishing.

Zecca, M. and Struhl, G. (2007a). Recruitment of cells into the Drosophila wing primordium by a feed-forward circuit of vestigial autoregulation. Development 134, 3001–3010.

Zecca, M. and Struhl, G. (2007b). Control of Drosophila wing growth by the vestigial quadrant enhancer. Development 134, 3011–3020.

Zecca, M., Basler, K. and Struhl, G. (1996). Direct and long-range action of a wingless morphogen gradient. Cell 87, 833–844.

